# Structural basis for differential targeting properties of small RNAs in plants and animals

**DOI:** 10.1101/2022.07.20.500788

**Authors:** Yao Xiao, Ian J. MacRae

## Abstract

microRNAs (miRNAs) regulate gene expression in plants and animals. Animals use miRNAs to sculpt the transcriptome, with each miRNA modestly repressing hundreds of otherwise unrelated targets. By contrast, each plant miRNA potently silences a small number of physiologically related targets. Here, we show that this major functional distinction depends on a minor structural difference between plant and animal Argonaute (AGO) proteins. A 7-amino-acid loop in the PIWI domain of *Arabidopsis* Argonaute10 (AtAGO10) reduces the affinity of the miRNA seed region for target RNAs. Swapping the PIWI-loop from human Argonaute2 (HsAGO2) into AtAGO10 increases seed strength and target-binding promiscuity, resembling animal miRNA targeting. Conversely, swapping the plant PIWI-loop into HsAGO2 increases targeting stringency and elevates target cleavage rates. The loop-swapped HsAGO2-siRNA complex silences targets more potently, with reduced off-targeting, than wild-type HsAGO2 and small interfering RNA (siRNA) duplexes in mammalian cells. Thus, tiny structural differences can tune the targeting properties of AGO proteins for distinct biological roles and HsAGO2 can be engineered for potential applications in siRNA therapeutics.

## Introduction

RNA silencing is a cellular process that down-regulates gene expression, wherein small RNAs, including microRNAs (miRNAs) and small interfering RNAs (siRNAs), function as guides for Argonaute (AGO) proteins. AGOs use small RNAs to identify complementary sites in transcripts targeted for repression. In many animals, and particularly vertebrates, AGO proteins are predominantly guided by miRNAs, an abundant class of small RNAs that are generated by the multistep processing of endogenous transcripts containing RNA hairpin structures (Ha & Kim, 2014). miRNA target recognition leads to a moderate (2-3 fold) reduction in gene expression via the recruitment of factors involved in translational repression and mRNA decay (Baek *et al*, 2008; Jonas & Izaurralde, 2015). The miRNA recognition elements (MREs) found in animal target mRNAs consist of a core segment of complementarity to miRNA guide (g) nucleotides g2–g7 (counting from the miRNA 5’ end), which is termed the miRNA seed region (Lewis *et al*, 2005). The short length (6-8 nt.) typical of animal MREs enables each miRNA to target hundreds of different mRNAs (Bartel, 2018), and miRNAs as a whole to regulate over half of protein-coding genes in mammals (Friedman *et al*, 2009). miRNAs are thus considered sculptors of the transcriptome in animals (Bartel, 2018).

Some animal AGOs can also be loaded with siRNAs, which are derived from long double-stranded RNAs or can be delivered as synthetic small RNA duplexes (Elbashir *et al*, 2001a; Elbashir *et al*, 2001b). siRNA-mediated silencing involves nearly complete guide-target base pairing, which elicits an endonuclease activity in many AGO proteins, particularly Argonaute2 (AGO2) in vertebrates (Liu *et al*, 2004; Meister *et al*, 2004; Rivas *et al*, 2005). Synthetic siRNAs are useful research tools for transient knock-down of target genes in cell culture and have emerged in the clinic as a novel approach to treating human disease (Syed, 2021; Weng *et al*, 2019). However, because vertebrate AGO2 evolved primarily for miRNA-mediated repression (Cheloufi *et al*, 2010; Chen *et al*, 2017; Cifuentes *et al*, 2010), siRNAs in humans inevitably also function as miRNAs and moderately repress hundreds of unintended targets, which can result in detrimental off-targeting and failure in clinical trials (Jackson & Linsley, 2010; Seok *et al*, 2018).

Plants produce miRNAs and several classes of siRNAs (Borges & Martienssen, 2015). In contrast to animals, miRNAs in plants are highly selective with an MRE core that is twice as long as typical mammalian MREs, comprising complementarity to g2–g13 (Allen *et al*, 2005). Plant miRNAs thereby restrict silencing to a small number of physiologically related targets (Jones-Rhoades & Bartel, 2004). The stringency of miRNA targeting in plants is explained, in part, by the observation that plant miRNAs often function through target RNA cleavage and thus, like siRNAs, require extensive target complementarity to act (Llave *et al*, 2002; Tang *et al*, 2003). However, plant miRNAs can also function through translational repression (Aukerman & Sakai, 2003; Brodersen *et al*, 2008; Chen, 2004; Gandikota *et al*, 2007; Li *et al*, 2013), and biochemical analysis of *Arabidopsis* AGO1 (AtAGO1) showed seed complementarity alone is insufficient for high-affinity target binding (Iwakawa & Tomari, 2013). Thus, plant AGO proteins appear to be inherently more selective in target recognition than AGOs in mammals.

We recently reported cryo-EM structures of *Arabidopsis* (AtAGO10) (Xiao *et al*, 2022). AtAGO10 is a close paralog of AtAGO1, with overlapping functions *in vivo* (Lynn *et al*, 1999), and similar biochemical activities *in vitro* (Ji *et al*, 2011). Surprisingly, despite the functional distinctions between plant and animal miRNAs, AtAGO10 structure is remarkably similar to human AGOs. Here, we explored the hypothesis that small structural differences between plant and animal AGOs are responsible for creating differential targeting properties. We identified a short loop in the PIWI domain of AtAGO10 that significantly influences targeting behavior. The loop is disordered in human AGO structures but forms a defined structure in AtAGO10. Structural analysis indicates the AtAGO10 loop decreases target affinity by stabilizing a closed AGO conformation that is incompatible with seed-pairing. Swapping the loop from HsAGO2 into AtAGO10, and vice versa produces an AtAGO10 variant with animal-like targeting properties and an HsAGO2 variant with plant-like targeting. Moreover, the loop-swapped HsAGO2-siRNA complex has superior on-target silencing potency and reduced off-targeting compared to conventional siRNAs and the wild-type HsAGO2-siRNA complex when introduced into mammalian cells. Combined results show that AGO-targeting behavior can be tuned by small changes and thus HsAGO2 can be engineered for improved therapeutic properties.

## Results

### Plant AGOs create a weak guide RNA seed region

AGO proteins create the seed region by pre-organizing guide nucleotides g2–g7 in a helical conformation, thereby lowering the entropic cost of initiating target binding (Parker *et al*, 2009; Schirle & MacRae, 2012; Wang *et al*, 2008). In animal AGOs, the affinity of the seed region for targets is quite high, with dissociation constants typically in the nM range, and seed-complementarity alone is sufficient for target recognition by miRNAs (Grimson *et al*, 2007; Lewis *et al*., 2005; Schirle *et al*, 2014; Wee *et al*, 2012). Curiously, although AtAGO10 pre-organizes the seed region in a fashion identical to human AGOs (Fig. 1A), AtAGO1, a close paralog of AtAGO10, was previously shown to not bind target RNAs with complementary limited to the seed (Iwakawa & Tomari, 2013). These observations indicate plant AGOs may dampen the affinity of the pre-organized seed through an unknown mechanism. To test this idea directly, we measured the affinity of purified the AtAGO10-miRNA complex for a seed-matched target RNA. Similar to results with AtAGO1 (Iwakawa & Tomari, 2013), AtAGO10 bound a seed-matched target with an 18-fold lower affinity than that of HsAGO2 using the same miRNA guide (Fig. 1B). Thus, although plant and animal AGOs pre-organize the seed region in the same way, plant AGOs possess a means of dampening seed strength.

**Fig 1.**
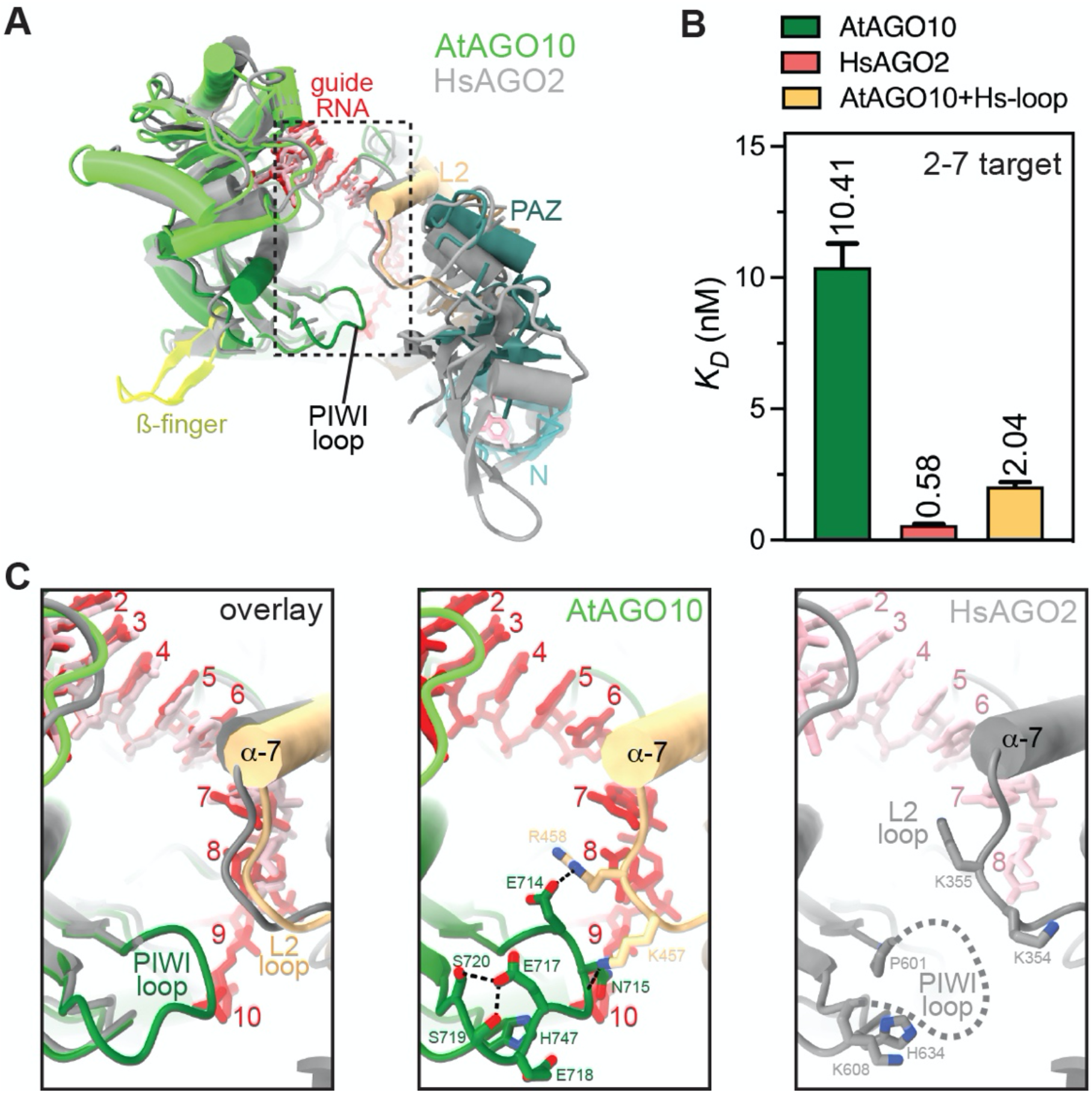
A structured PIWI domain loop distinguishes AtAGO10 from HsAGO2. **A**. Superposition of AtAGO10 (colored) and HsAGO2 (gray) bound to guide RNAs (red/pink). PIWI-loop indicated. **B**. Dissociation constant values for wild-type AtAGO10, HsAGO2, and the AtAGO10+Hs-loop mutant binding to a target RNA with base-pairing complementarity to the miRNA seed region (guide nucleotides 2–7). **C**. Close-up views of boxed region in panel A showing the PIWI-loop is ordered in AtAGO10 and disordered in HsAGO2.

### A loop in the AtAGO10 PIWI domain dampens seed pairing

To understand how AtAGO10 dampens its seed region, we compared the AtAGO10-miRNA structure to human AGO-miRNA structures. There are multiple amino acid substitutions surrounding the seed region, but the largest difference, in terms of protein backbone conformation, is a PIWI domain loop immediately downstream of the seed (Fig. 1C). This loop is disordered in HsAGO2 crystal structures but adopts a defined conformation in AtAGO10 (Fig. 1C, S1A). The AtAGO10 PIWI loop contacts the L2 domain adjacent to helix-7, which modulates the kinetics of seed-pairing in HsAGO2 (Klum *et al*, 2017). Thus, the PIWI loop was a likely candidate for dampening the plant AGO seed.

To test this idea, we created an AtAGO10 mutant (AtAGO10+Hs-loop) in which the endogenous PIWI loop (residues 713–720) was replaced by the corresponding loop sequence from HsAGO2 (residues 602–608). The AtAGO10+Hs-loop mutant bound a seed-matched target with a 5-fold greater affinity than wild-type AtAGO10 (Fig. 1B). Residues involved in stabilizing the PIWI loop conformation are largely conserved in AtAGO1 (Fig. S1B) and swapping the PIWI loop sequence of AtAGO1 into AtAGO10 had only modest effects on target affinity (Fig. S1C). Thus, AtAGO1 and AtAGO10 have stabilized PIWI loops that dampen seed pairing compared to HsAGO2.

### The PIWI loop increases the dynamic range of target recognition by AtAGO10

To understand how the PIWI loop affects miRNA-target pairing beyond the seed, we measured the affinities of wild-type AtAGO10 and AtAGO10+Hs-loop for a series of target RNAs with increasing miRNA complementarity (Fig. 2A). Target affinity of AtAGO10 varied 33-fold between targets with complementarity to g2–g8 (*K*_*D*_ = 1.33 ± 0.14 nM) and g2–g21 (*K*_*D*_ = 0.040 ± 0.007 nM) (Fig. 2B,C). Target affinities of AtAGO10 bearing the PIWI loop of AtAGO1 were similar (Fig. S1D, E). By contrast, AtAGO10+Hs-loop target affinities varied only 4-fold between targets with complementarity to g2–g8 (0.38 ± 0.03 nM) and g2–g21 (0.095 ± 0.012 nM) (Fig. 2B,C). Thus, by weakening the strength of seed pairing, the structured PIWI loop effectively expands the dynamic range of miRNA target recognition by AtAGO10.

**Fig. 2.**
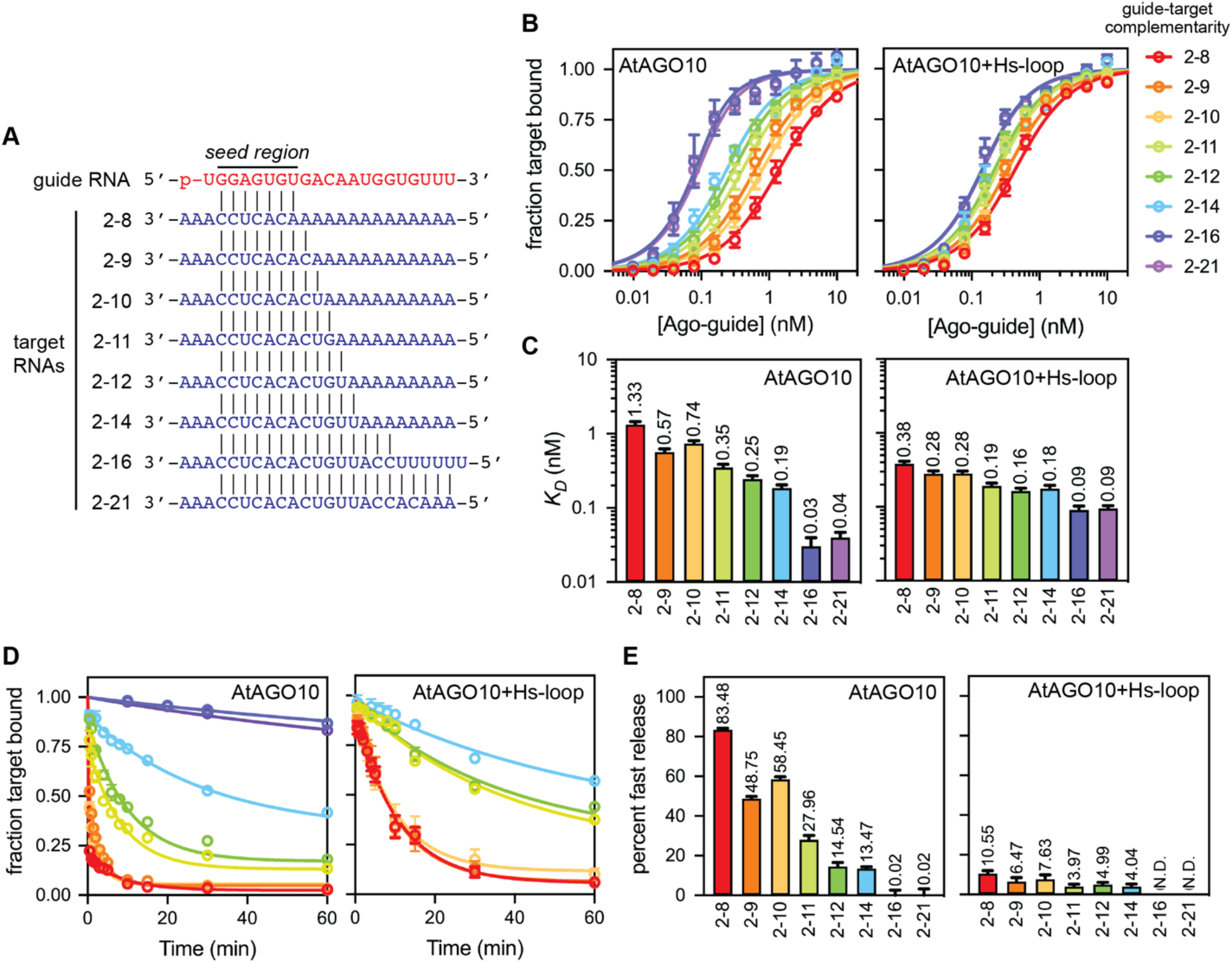
The AtAGO10 PIWI loop contributes to creating the extended MREs observed in plants. **A**. Base-pairing schematic of guide RNA and target RNAs used in panels B-E and Fig. 4. **B**. Fraction target RNA bound versus [AGO-guide] for wild-type AtAGO10 and the AtAGO10+Hs-loop mutant. **C**. Dissociation constants derived from data in panel B. **D**. Fraction of ^32^P-labeled target RNAs bound by AtAGO10 or AtAGO10+Hs-loop after the addition of excess unlabeled target RNA versus time. Data were fit with a two-phase decay with a fixed *k*_*off,fast*_ value for 4.6 min^-1^ for all curve fits. **E**. Fraction of target RNA released in the fast phase of curve fits in panel D. Data points represent the mean value of three experimental replicates. Error bars indicate standard deviation.

### The AtAGO10 PIWI loop connects stable seed pairing to downstream miRNA-target interactions

We also measured target release rates from wild-type AtAGO10 and AtAGO10+Hs-loop (Fig. 2D). The release of a target with complementarity to the extended seed region (2–8 target) from AtAGO10 followed a two-phase exponential, with ∼85% of the target molecules released rapidly (*k*_*off,fast*_ = 4.6 min^-1^) and ∼15% of the target molecules released 40-times more slowly (*k*_*off,slow*_ = 0.12 min^-1^) (Fig. 2D). The observed biphasic kinetics indicate that wild-type AtAGO10 engages the 2–8 target in at least two distinct binding modes. By contrast, the release of the 2–8 target from AtAGO10+Hs-loop was well described by a one-phase exponential, indicating a single binding mode. Notably, the rate of 2–8 target release from AtAGO10+Hs-loop (0.13 min^-1^) closely matches the *k*_*off,slow*_ rate observed in the bi-phasic target release from wild-type AtAGO10 (Fig. 2D). These results suggest that a minor fraction (∼15%) of 2–8 target molecules fully engage the seed region when bound to wild-type AtAGO10 while the majority (∼85%) of target molecules do not. The fraction of target molecules released from AtAGO10 in the fast phase decreased with increasing guide-target complementarity (Fig. 2E). Thus, for most binding events, the structured PIWI loop causes AtAGO10 to rapidly release seed-matched targets in the absence of downstream miRNA-target pairing.

### Structural model for miRNA targeting selectivity in plants

We propose that the difference in MRE size between plants and animals arises, to a large extent, from differences in plant and animal PIWI loop structures. In this model, the stabilized AtAgo10 PIWI loop reaches across the central RNA-binding cleft to contact a loop in the L2 domain (Fig. 1C, S1A). This connection may dampen seed pairing by stabilizing the closed position of helix-7, which is directly tethered to the L2 loop and pivots between open and closed positions to facilitate the making and breaking of miRNA-target base pairs in the 3’ half of the seed (Klum *et al*., 2017; Schirle *et al*., 2014). By contrast, the PIWI loop in HsAgo2 is unstructured and thus cannot influence helix-7 (Fig. 1C, S1A). We expect miRNA-target pairing downstream of the seed physically disrupts the connection between PIWI and L2 loops, reducing the propensity of helix-7 to occupy its closed, seed-breaking position. The AtAgo10 PIWI loop thus makes stable seed pairing dependent on downstream miRNA-target interactions and thereby creates the extended MREs observed in plants (Allen *et al*., 2005).

### The AtAGO10 PIWI loop adopts a structured conformation when swapped into HsAGO2

We next asked if the AtAGO10 PIWI loop can also adopt a stabilized conformation when swapped into HsAGO2. We produced an HsAGO2 mutant (HsAGO2+At-loop) in which the endogenous PIWI loop sequence (residues 602–608) was replaced by the corresponding PIWI loop sequence of AtAGO10 (residues 713–720). We then determined the 2.5 Å resolution crystal structure of the HsAGO2+At-loop mutant bound to the miRNA miR-122 (Fig 3A, Table S1). The structure reveals that the swapped PIWI loop adopts a defined conformation (Fig. 3B), likely stabilized by multiple internal hydrogen bonds within the loop: the sidechains of E602 and N603, the N603 mainchain carbonyl and E605 sidechain, and the sidechains of E605 and S608 are within hydrogen-bonding distances of each other. The stabilized loop reaches across the central cleft and forms two interactions with the L2 loop: the N603 sidechain is within hydrogen-bonding distance of the mainchain amine of K355, and the mainchain carbonyl of G604 may form a hydrogen bond with the sidechain of K354.

**Fig. 3.**
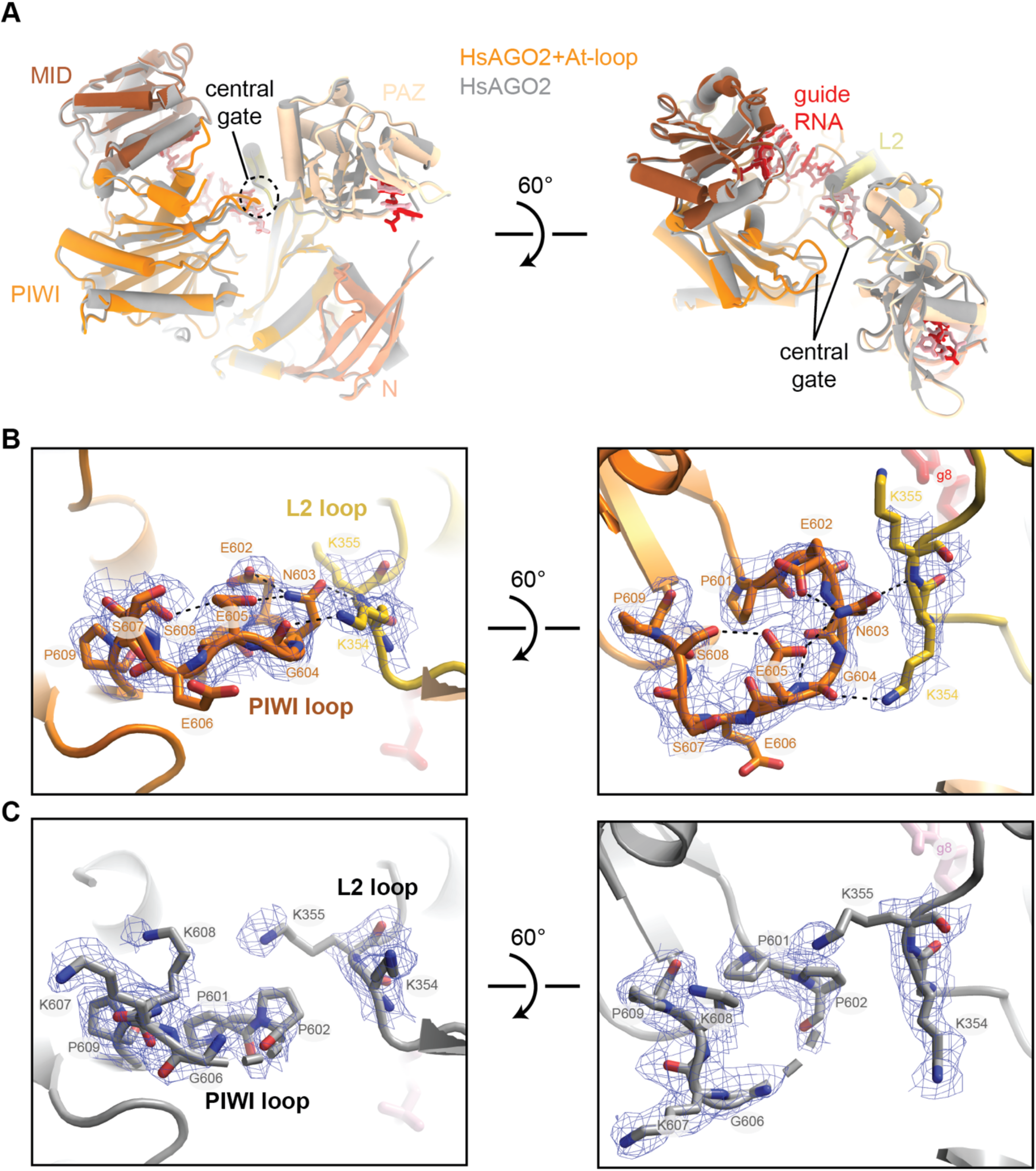
The AtAGO10 PIWI loop adopts a stable conformation when swapped into HsAGO2. **A**. Superposition of wild-type HsAGO2 (gray) and the HsAGO2+At-loop mutant (colored) crystal structures reveals nearly identical structures. **B**. Close-up view of PIWI loop in the HsAGO2+At-loop mutant crystal structure. Blue mesh shows the refined electron density map. **C**. Close-up view of PIWI loop in the wild-type HsAGO2 crystal structure. Dashes indicate disordered residues in the PIWI-loop. Blue mesh shows the refined electron density map.

For direct comparison, we crystallized and determined the 2.5 Å resolution structure of wild-type HsAGO2 bound to miR-122 side-by-side with the mutant complex. In contrast to the HsAGO2+At-loop mutant, there is no contiguous electron density corresponding to the center of the wild-type HsAGO2 PIWI loop (residues 603-606 are disordered), and no interactions between the PIWI loop and L2 loop are observed (Fig. 3C). Except for the difference in the PIWI loops, the wild-type HsAGO2 and HsAGO2+At-loop mutant structures are nearly identical, with an RMSD of 0.61 Å for all equivalent Cα atoms (Fig. 3A). Thus, the swapped-in AtAGO10 PIWI loop adopts a defined conformation within HsAGO2 without otherwise impacting the overall structure.

### HsAGO2+At-loop has increased targeting stringency compared to HsAGO2

We compared the target affinities of the HsAGO2+At-loop mutant to those of wild-type HsAGO2. Consistent with the hypothesis that a stabilized PIWI loop dampens seed-pairing, HsAGO2+At-loop bound the 2–8 target with ∼75-fold lower affinity than wild-type HsAGO2 (*K*_*D*_ = 6.43±.24 and 0.086±0.005 nM for HsAGO2+At-loop and wild-type, respectively) (Fig. 4A, B). By contrast, HsAGO2+At-loop and wild-type HsAGO2 bound the 2–21 complementarity target with comparable affinities (*K*_*D*_ = 5±1 pM and *K*_*D*_ = 10±1 pM for mutant and wild-type, respectively) (Fig. 4A, B). Notably, target affinities of HsAGO2+At-loop varied over 1000-fold between the 2–8 target to the 2–21 target. This dynamic range is two orders of magnitude greater than the 9-fold difference observed for the corresponding target affinities of wild-type HsAGO2. Thus, HsAGO2+At-loop mutant recognizes targets with extensive guide-complementary targets as well as wild-type HsAGO2 but is far more discerning against targets with limited complementarity.

**Fig. 4.**
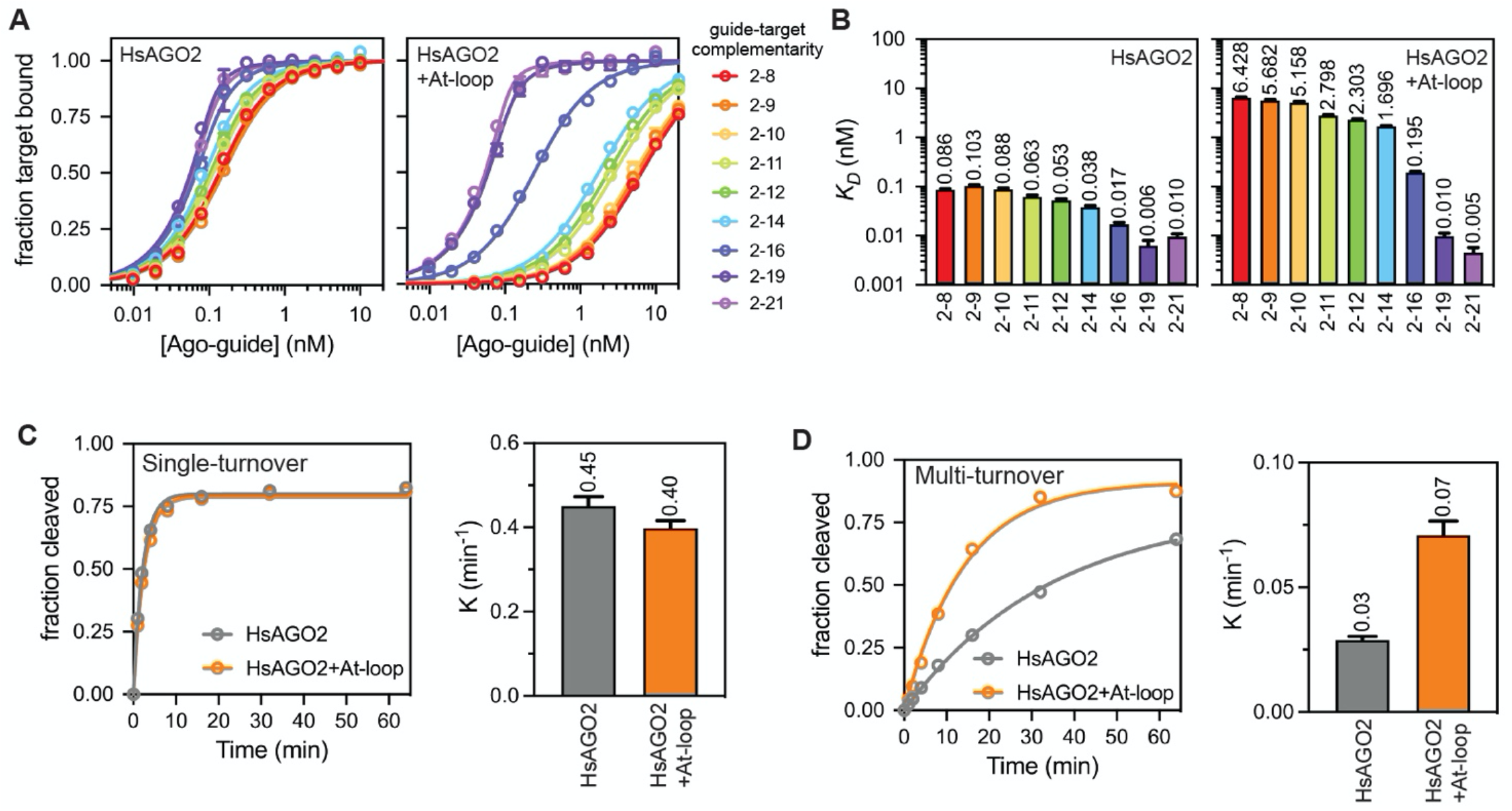
Swapping the AtAGO10 PIWI loop into HsAGO2 increases targeting selectivity and catalytic turnover. **A**. Fraction target RNA bound versus [AGO-guide] for wild-type HsAGO2 and the HsAGO2+At-loop mutant. Guide and target RNAs used are the same as shown in Fig. 2A. **B**. Dissociation constants derived from data in panel A. **C**. Fraction of a 2–21 complementary target RNA cleaved by HsAGO2 and HsAGO2+At-loop under single turnover conditions versus time (left panel). The right panel shows first-order rate constants for the single-turnover condition. **D**. Fraction of a 2–21 complementary target RNA cleaved by HsAGO2 and HsAGO2+At-loop under multiple turnover conditions versus time (left panel). The right panel shows first-order rate constants for the multiple-turnover condition.

### HsAGO2+At-loop has a higher multiple turnover cleavage rate than HsAGO2

We next compared target RNA cleavage rates by wild-type HsAGO2 and the HsAGO2+At-loop mutant. Under single-turnover conditions ([AGO2-guide complex] = 5nM, [target RNA] = 1nM), wild-type HsAGO2 and HsAGO2+At-loop cleaved the 2–21 complementary target at the same rate (Fig. 4C). Both wild-type and mutant also behaved the same with respect to 2–19 and 2–16 complementary targets under single-turnover conditions (Fig. S2A,B). Thus, the PIWI loop does not impact *k*_*cat*_, the rate constant describing hydrolysis of bound target RNAs. By contrast, under multiple turnover conditions ([AGO2-guide complex] = 1nM, [target] = 5nM), HsAGO2+At-loop shows a significantly higher target cleavage rate than HsAGO2 (Fig. 4E,F). HsAGO2+At-loop also displayed increased multiple-turnover cleavage rates for 2–19 and 2–16 complementary targets (Fig. S2C,D) Increased multiple turnover rates are likely due to accelerated target release as the At-loop reduces the affinity of HsAGO for the 2-10 target (Fig. 4B), which mimics the 3’ cleavage product, as previous results showed multiple turnover is dictated by product release (Salomon *et al*, 2015; Wee *et al*., 2012). The combined results show that swapping in the AtAGO10 PIWI loop improves both substrate-binding selectivity and turnover kinetics of target RNAs by HsAGO2.

### Swapping the AtAGO10 PIWI loop into HsAGO2 improves siRNA-mediated silencing in mammalian cells

After finding that HsAGO2+At-loop has increased target selectivity and catalytic turnover *in vitro*, we wondered if the engineered HsAGO2 has enhanced siRNA-mediated silencing properties in mammalian cells. We transfected various amounts of the purified HsAGO2+At-loop-guide-RNA complex into HEK293 cells along with luciferase reporters bearing complementary sites to either g2–g8 (miRNA-target) or g2–g21 (siRNA-target) of the guide RNA (Fig. 5A). Our rationale was that the miRNA-target reports on miRNA-mediated silencing (off-targeting) by the complex and that the siRNA-target reports siRNA-mediated (on-target) silencing. For comparison, we also transfected HEK293 cells with the luciferase reporters and either the wild-type HsAGO2-guide-RNA complex or an equivalent amount of siRNA duplex.

**Fig. 5.**
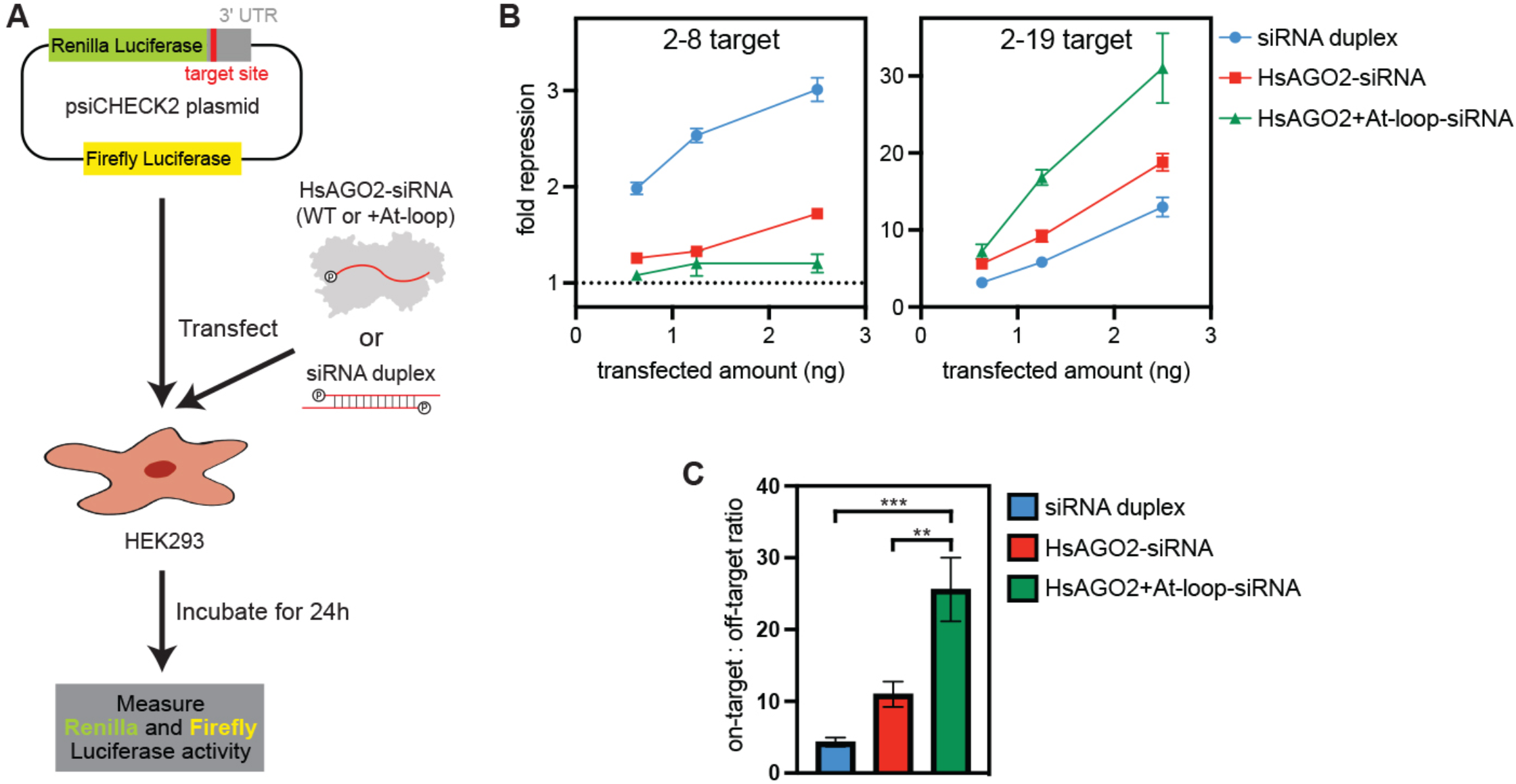
HsAGO2+At-loop has improved RNAi selectivity and potency in mammalian cells. **A**. Schematic of RNAi experiment. HEK293 cells were transfected with a plasmid encoding *Renilla* luciferase with a single target site and targeting wild-type or +At-loop mutant HsAGO2-siRNA complexes, or siRNA duplexes. **B**. Fold repression of *Renilla* expression (compared to reporter plasmid transfection alone) versus the amount of transfected siRNA duplex, wild-type HsAGO2-siRNA complex, or HsAGO2+At-loop-siRNA complex. The 2–8 target (left panel) indicates off-targeting. The 2–19 target (right panel) indicates on-targeting. **C**. Ratio of on-targeting to off-targeting at the highest transfected amounts in panel B. Unpaired t-test p values: ***p = 0.0002, **p = 0.0061.

The engineered HsAGO2+At-loop-guide-RNA complex had minor effects on the expression of the miRNA-target reporter, repressing luciferase levels 1.1–1.2-fold relative to transfection of the reporter alone (Fig. 5B). Wild-type HsAGO2-guide-RNA and the siRNA duplex both displayed stronger off-targeting, repressing the miRNA-target 1.3–1.7-fold and 2.1–3.3-fold, respectively. By contrast, HsAGO2+At-loop effectively silenced the siRNA-target reporter, reducing expression 7–31-fold (Fig. 5B). Wild-type HsAGO2-guide-RNA and the siRNA duplex were both less potent, repressing the siRNA-target 5–19-fold and 3–13-fold, respectively (Fig. 5B). Thus, consistent with *in vitro* measurements, the engineered HsAGO2+At-loop displays superior RNAi properties, with enhanced on-target silencing and reduced off-targeting in mammalian cells (Fig. 5C).

## Discussion

Argonaute proteins have long been known to reshape the fundamental properties of RNA:RNA hybridization for guide-target pairing (Wee *et al*., 2012). This feature enables AGOs to conduct rapid and accurate target searches (Chandradoss *et al*, 2015; Salomon *et al*., 2015). Based on our results and the previous work of others (Becker *et al*, 2019; Iwakawa & Tomari, 2013; Ober-Reynolds *et al*, 2022; Wee *et al*., 2012), we suggest that controlling guide-target hybridization also allows different AGOs to adapt distinct small RNA classes to discrete biological roles. We show that the identity of a small PIWI domain loop can have dramatic effects on guide-target hybridization in the miRNA seed, giving rise to the distinct MREs found in plants and animals. Notably, the PIWI loop does not directly contact the guide or target nucleotides involved in seed-pairing (Fig. 1). We, therefore, suggest that reshaping guide-target hybridization involves controlling the dynamics of AGO conformational changes in helix-7 that are known to be necessary for seed-pairing (Klum *et al*., 2017; Schirle *et al*., 2014). Notably, although swapping the human PIWI loop into AtAGO10 significantly increased the affinity for a seed-complementary target, the overall affinity was not as high as the affinity of HsAGO2 for the same target RNA (Fig. 1B). Similarly, swapping the plant PIWI loop into HsAGO2 decreased affinity for an extended seed target to a level 5-fold lower than the affinity of AtAGO10 for the same target RNA (2-8 target in Fig. 1B, 4B). Thus, additional structural differences, beyond the PIWI loop, contribute to the distinct target-recognition properties of AtAGO10 and HsAGO2.

Conventional RNAi in both laboratory and clinical settings is usually induced via the introduction of siRNA duplexes into target cells or tissues. This approach requires the loading of siRNAs into endogenous HsAGO2 molecules upon reaching the cytoplasm as the inhibitory effect of siRNAs is determined by the abundance of the HsAGO2-siRNA complex (Wei *et al*, 2011). Circumventing this requirement, recent studies have shown that the direct introduction of preassembled HsAGO2-siRNA complexes can induce the knock-down target mRNAs in *Cryptosporidium* parasites wherein endogenous AGOs are not present (Castellanos-Gonzalez *et al*, 2016). Similarly, preassembled HsAGO2-siRNA complexes were shown to potently knock-down target gene expression in human 293T cells in culture and in B16-F10 tumors in mice (Castellanos-Gonzalez *et al*., 2016). Our results indicate that this approach may be further advanced through the development of HsAGO-siRNA complexes with enhanced gene-silencing properties. A comparison of target RNA cleavage by HsAGO2 and *Drosophila* AGO2 revealed that the fly protein, which evolved for siRNA-mediated silencing, cleaves targets at a rate that is orders of magnitude faster than the human enzyme, which is evolved for miRNA-mediated repression (Wee *et al*., 2012). Taken with our finding that the simple replacement 7 amino acids significantly alters targeting behavior, we envision further improvements in the catalytic activity of HsAGO2 are possible through additional minor adjustments. Therefore, we suggest incorporating the AtAGO10 PIWI loop, as well as any other small changes that impact conformational dynamics, may result in an HsAGO2-siRNA complex with significantly enhanced siRNA-directed silencing potency and negligible off-targeting.

## Acknowledgments

We are grateful to MacRae lab members for their advice. The research was funded by NIH grant R35GM127090 to I.J.M.

## Supplementary Materials

### Methods and Materials

#### Bacterial strains and plasmids

Bacteria used for cloning were chemically competent *E. coli* OmniMAX™ (Thermo Fisher). Bacteria used for the production of bacmid DNAs were DH10Bac™ chemically competent *E. coli* (Thermo Fisher).

#### Bacterial media and growth conditions

All bacterial cultures were grown in Luria-Bertani (LB) medium at 37 °C. When needed, media was supplemented with one or more of the following antibiotics at the following concentrations: ampicillin (100 μg/mL), kanamycin (40 μg/mL), tetracycline (5 μg/mL), gentamycin (7 μg/mL), 5-Bromo-4-Chloro-3-Indolyl β-D-Galactopyranoside (X-gal, 25 μg/mL in dimethylformamide), and/or Isopropyl β-D-1-thiogalactopyranoside (IPTG, 1 mM).

#### Insect cell media and growth conditions

Sf9 cells were grown in Lonza Insect XPRESS™ medium supplemented with 1x Gibco™ Antibiotic-Antimycotic in suspension at 27 °C.

#### Cloning and mutagenesis

The Piwi-loop mutated AtAGO10/HsAGO2 were generated by first amplifying two AtAGO10/HsAGO2 fragments with PCR primers overlapping at codons replacing the PIWI loop, and then assembling two gel purified PCR fragments with digested pFastBac HTA by NEBuilder® HiFi DNA Assembly Master Mix.

#### Preparation of AtAgo10-guide RNA complex

His_6_-Flag-Tev-tagged AtAgo10 proteins were expressed in Sf9 cells using a baculovirus system (Invitrogen). 60 h cultured 750 mL 3.4 × 10^6^ cells/mL Sf9 cells infected with ∼15 mL virus were harvested by centrifugation. Usually, two of these cell pellets were combined and suspended in ∼200mL Lysis Buffer (50 mM NaH2PO4, pH 8, 300 mM NaCl, 5% glycerol, 20 mM imidazole, 0.5 mM TCEP). Resuspended cells were lysed by passing twice pass through a M-110P lab homogenizer (Microfluidics). The resulting total cell lysate was clarified by centrifugation (30,000 x g for 25 min) and the soluble fraction was applied to 8 mL packed Ni-NTA resin (Qiagen) and gently rocked at 4°C for 1.5 hours in 50 mL conical tubes. Resin was pelleted by brief centrifugation and the supernatant solution was discarded. The resin was washed with ∼50 mL ice cold Nickel Wash Buffer (300 mM NaCl, 20 mM imidazole, 0.5 mM TCEP, 50 mM Tris, pH 8). Centrifugation/wash steps were repeated a total of three times. Co-purified cellular RNAs were degraded by incubating with 400U micrococcal nuclease (Clontech) on-resin in ∼25 mL of Nickel Wash Buffer supplemented with 5 mM CaCl2 at room temperature for ∼1h. The nuclease-treated resin was washed three times again with Nickel Wash Buffer and then eluted in four column volumes of Nickel Elution Buffer (300 mM NaCl, 300 mM imidazole, 0.5 mM TCEP, 50 mM Tris, pH 8). Eluted protein was incubated with 24 nmol synthetic guide RNA and 150 µg TEV protease during ∼2h dialysis against 1 L of Dialysis Buffer (300 mM NaCl, 0.5 mM TCEP, 50 mM Tris, pH 8) at 4°C. After dialysis, the capture resin was prepared by incubating 345.6 uL packed High Capacity Neutravidin Resin (Thermo Fisher) with 28.8 nmol capture oligo in wash A buffer (100 mM KOAc, 2 mM MgOAc, 0.01% CHAPS, 30 mM Tris, pH 8) for 30 min at 4 °C, following by 10 mL wash A buffer wash. Dialyzed protein supplemented with 0.01% CHAPS and 2 mM MgOAc was then incubated with capture resin at RT for 1 hour (without rocking!!!! Just gently inverting the tube every 5-10 minutes). Then, the resin was washed by three times with 10 mL Wash A followed by three times with 10 mL Wash B (2 M KOAc, 2 mM MgOAc, 0.01% CHAPS, 30 mM Tris, pH 8) and Wash C (1 M KOAc, 2 mM MgOAc, 0.01% CHAPS, 30 mM Tris, pH 8). The resin was then re-suspended in 1900 µL Wash C with 57.6 nmol competitor DNA at RT for ∼2 hours (Gently inverting the tube every 5-10min). The elute was dialyzed in 1 L Q dialyzing buffer (150 mM NaCl, 0.01% CHAPS, 0.5 mM TCEP, 20 mM Tris, pH 8) at 4 °C overnight. After dialysis, 240 µl Q Sepharose Fast Flow anion exchange resin slurry (GE Healthcare) was equilibrated in Q dialyzing buffer. The dialyzed protein was then passed through this resin to remove unbound oligonucleotides, and flow-through solution was collected. The flow-through was then concentrated while buffer exchanging to 1∼3 mg/mL in Tris Crystal buffer (10 mM Tris pH 8, 100 mM NaCl, 0.5 mM TCEP). The concentrated protein was aliquoted, flash frozen in liquid N_2_ and stored at -80 °C. Concentration of the AtAgo10-guide RNA complex was determined by Bradford assay with BSA as a standard.

#### Preparation of HsAgo2-guide RNA complex

His_6_-Flag-Tev-tagged HsAgo2 proteins were expressed in Sf9 cells using a baculovirus system (Invitrogen). 60 h cultured 750 mL 3.4 × 10^6^ cells/mL Sf9 cells infected with ∼15 mL virus were harvested by centrifugation. One pellet was suspended in ∼100 mL Lysis Buffer (50 mM NaH2PO4, pH 8, 300 mM NaCl, 5% glycerol, 20 mM imidazole, 0.5 mM TCEP). Resuspended cells were lysed by passing twice pass through a M-110P lab homogenizer (Microfluidics). The resulting total cell lysate was clarified by centrifugation (30,000 x g for 25 min) and the soluble fraction was applied to 3 mL packed Ni-NTA resin (Qiagen) and gently rocked at 4°C for 1.5 hours in 50 mL conical tubes. Resin was pelleted by brief centrifugation and the supernatant solution was discarded. The resin was washed with ∼50 mL ice cold Nickel Wash Buffer (300 mM NaCl, 20 mM imidazole, 0.5 mM TCEP, 50 mM Tris, pH 8). Centrifugation/wash steps were repeated a total of three times. Co-purified cellular RNAs were degraded by incubating with 200U micrococcal nuclease (Clontech) on-resin in ∼25 mL of Nickel Wash Buffer supplemented with 5 mM CaCl2 at room temperature for ∼1h. The nuclease-treated resin was washed three times again with Nickel Wash Buffer and then eluted in four column volumes of Nickel Elution Buffer (300 mM NaCl, 300 mM imidazole, 0.5 mM TCEP, 50 mM Tris, pH 8). Eluted protein was incubated with 24 nmol synthetic guide RNA and 150 µg TEV protease during an overnight dialysis against 1∼2 liters of Dialysis Buffer (300 mM NaCl, 0.5 mM TCEP, 50 mM Tris, pH 8) at 4°C. After dialysis, the capture resin was prepared by incubating 345.6 μL packed High Capacity Neutravidin Resin (Thermo Fisher) with 28.8 nmol capture oligo in wash A buffer (100 mM KOAc, 2 mM MgOAc, 0.01% CHAPS, 30 mM Tris, pH 8) for 30 min at 4 °C, following by 10 mL wash A buffer wash. Dialyzed protein supplemented with 0.01% CHAPS and 2 mM MgOAc was then incubated with capture resin at RT for 1 hour (without rocking!!!! Just gently inverting the tube every 5-10 minutes). Then, the resin was washed by three times with 10 mL Wash A followed by three times with 10 ml Wash B (2 M KOAc, 2 mM MgOAc, 0.01% CHAPS, 30 mM Tris, pH 8) and Wash C (1 M KOAc, 2 mM MgOAc, 0.01% CHAPS, 30 mM Tris, pH 8). The resin was then re-suspended in 1900 µL Wash C with 57.6 nmol competitor DNA at RT for ∼2 hours (Gently inverting the tube every 5-10min). The elute was dialyzed in 1 L Q dialyzing buffer (150 mM NaCl, 0.01% CHAPS, 0.5 mM TCEP, 20 mM Tris, pH 8) at 4 °C for 1 hour, then moved to fresh 1 liter Q dialyzing buffer for another 2 hours. When the dialyzing was near completion, 240 µl Q Sepharose Fast Flow anion exchange resin slurry (GE Healthcare) was equilibrated in Q dialyzing buffer. The dialyzed protein was then passed through this resin to remove unbound oligonucleotides, and flow-through solution was collected. The flow-through was then concentrated while buffer exchanging to 1∼3 mg/mL in Tris Crystal buffer (10 mM Tris pH 8, 100 mM NaCl, 0.5 mM TCEP). The concentrated protein was aliquoted, flash frozen in liquid N_2_ and stored at -80 °C. Concentration of the HsAgo2-guide RNA complex was determined by Bradford assay with BSA as a standard.

#### Equilibrium target binding assays

Equilibrium dissociation constants were determined as described previously (Schirle et al., 2014). Briefly, various concentrations of the AtAgo10_D795A-guide RNA, AtAgo10+Hs-loop_D795A-guide RNA, HsAGO2-guide RNA or HsAGO2+At-loop-guide RNA samples were incubated with 0.1 nM ^32^P 5’-radiolabeled target RNA in binding reaction buffer (30 mM Tris pH 8, 100 mM potassium acetate, 2 mM Magnesium acetate, 0.5 mM TCEP, 0.005% (v/v) NP-40, 0.01 mg/mL baker’s yeast tRNA), in a reaction volume of 100 μL at room temperature for 60 min. Using a dot-blot apparatus (GE Healthcare Life Sciences), protein-RNA complexes were captured on Protran nitrocellulose membrane (0.45 μm pore size, Whatman, GE Healthcare Life Sciences) and unbound RNA on Hybond Nylon membrane (Amersham, GE Healthcare Life Sciences). Samples were applied with vacuum and immediately washed once with 100 μL of ice-cold wash buffer (30 mM Tris pH 8.0, 100 mM potassium acetate, 2 mM Magnesium acetate, 0.5 mM TCEP). Membranes were air-dried and 32P signal was visualized by phosphor-imaging. ImageQuant (GE Healthcare Life Sciences) was used to quantify data and dissociation constants calculated using Prism version 6.0g (GraphPad Software, Inc.), using the following formula, which accounts for potential ligand depletion (Wee et al., 2012):

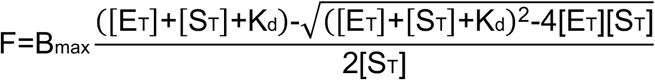

where F = fraction of target bound, B_max_ = calculated maximum number of binding sites, [E_T_] = total enzyme concentration, [S_T_] = total target concentration, and K_d_ = apparent equilibrium dissociation constant.

#### Target dissociation assay

Target dissociation rates were determined by incubating AtAgo10_D795A-guide RNA, AtAgo10+Hs-loop_D795A-guide RNA, HsAGO2-guide RNA or HsAGO2+At-loop-guide RNA samples with 0.1 nM ^32^P 5’-radiolabelled target RNA in binding reaction buffer (30 mM Tris pH 8.0, 100 mM potassium acetate, 2 mM Magnesium acetate, 0.5 mM TCEP, 0.005% (v/v) NP-40, 0.01 mg/mL baker’s yeast tRNA) in a single reaction with a volume of 100 μL per time point planned for the experiment (for example, 1,000 µl for 10 time points) at room temperature for 60 min. The concentration of protein complex was 5nM for 2-8 target or 2.5nM for all other targets measured. After sample equilibration, a zero-time point was taken by applying 100 µl of the reaction to the dot-blot apparatus under vacuum, followed by 100 μL of ice-cold wash buffer (30 mM Tris pH 8.0, 100 mM potassium acetate, 2 mM Magnesium acetate, 0.5 mM TCEP). The dissociation time course was started by the addition of 300 nM (final concentration) unlabelled target RNA. Aliquots of 100 μL were taken at various times and immediately applied to a dot-blot apparatus under vacuum, followed by 100 μL of ice-cold wash buffer. Time points ranged from 0.25 to 100 min. Membranes were air-dried and visualized by phosphorimaging. Quantification of the 32P signal was performed using ImageQuant TL (GE Healthcare). The fraction of target RNA bound was calculated as the ratio of bound to total (bound + free) target RNA for various concentrations of AGO-guide complexes. Dissociation rates were calculated by plotting data as fraction bound versus time and fitting to a two-phase decay curve with shared Kfast value using Prism v.8.0 (GraphPad).

#### Target Cleavage assay

For single turnover slicing assay, purified HsAGO2-guide RNA or HsAGO2+At-loop-guide RNA complex (10 nM, final concentration) were incubated at 37c with complementary ^32^P 5’-radiolabelled target RNAs (2 nM, final concentrations) in reaction buffer composed of 30 mM Tris pH 8.0, 100 mM potassium acetate, 2 mM Magnesium acetate, 0.5 mM TCEP, and 0.01 mg/mL baker’s yeast tRNA. For multiple turnover slicing, purified HsAGO2-guide RNA complexes or HsAGO2+At-loop-guide RNA complex (1 nM, final concentration) were incubated at 37c with complementary ^32^P 5’-radiolabelled target RNAs (5 nM, final concentrations) in reaction buffer. Target cleavage was stopped at various times by mixing aliquots of each reaction with an equal volume of denaturing gel loading buffer (98% w/v formamide, 0.025% xylene cyanol, 0.025% w/v bromophenol blue, 10 mM EDTA pH 8.0). Intact and cleaved target RNAs were resolved by denaturing PAGE (15%) and visualized by phosphorimaging. Quantification of signal was performed using ImageQuant TL (GE Healthcare).

#### Crystallization and diffraction data collection

HsAGO2-miR122 and HsAGO2+At-loop-miR122 crystals were grown by hanging drop vapor diffusion at 20°C and appeared in 24 hours. Drops contained a 1.0:0.8 ratio of protein (1 mg/mL) to reservoir solution (0.1M Tris pH8, 10mM MgCl2, 0.1M Phenol, 15% PEG3350, 10% Isopropanol). After growing for one week, crystals were harvested for x-ray data collection by soaking first in reservoir solution containing 25% ethylene glycol as cryo-protectant. Following cryo-protection, crystals were cryo-cooled by plunging into liquid N2. Data were collected on beam lines 9-2 at the Stanford Synchrotron Radiation Lightsource (SSRL). Data were processed using XDS and Aimless (Kabsch, 2010; Winn *et al*, 2011).

#### Model building and refinement

The HsAGO2-miR122 and HsAGO2+At-loop-miR122 structures were solved by molecular replacement using a previously determined guide-only structure (PDB:4OLA) as search models with Phaser-MR in the PHENIX graphical user interface (Adams *et al*, 2010). Models were built using Coot (Emsley *et al*, 2010) and were submitted to XYZ coordinate, Occupancies, and B-factor refinement using PHENIX. Model building and refinement continued iteratively until all interpretable electron density was modeled. Water molecules were updated automatically in PHENIX. All structure figures were generated with PyMOL (Schrödinger, LLC).

#### Luciferase assay

Target sites were cloned into psiCHECK2 plasmid (Promega Corporation) using Xho1 and Not1 sites. The siRNA duplex was made by denaturing 1:1 mixture of guide and passenger strands in annealing buffer (10 mM Tris, pH8.0, 50 mM NaCl) at 95°C for 2 min, then cooling to room temperature by placing on the benchtop. Luciferase reporter assays were performed with HEK293 cells. The HEK293 cells were plate at 10^4^ cells with 80ul DMEM complete media per well in 96-well white plate (Thermal Scientific, Cat#136101) and cultured in a temperature-, CO^2^-, and humidity-controlled cell incubator for ∼ 4 h to allow cells attach to the bottom of the plate. Cells in each well were first transfected with 40 ng psiCHECK2 plasmid using Lipofectamine 2000 (Invitrogen) based on manufacturer’s protocol. Then, 2.5μL 2.5pmol siRNA duplex, HsAGO2-siRNA or HsAGO2+At-loop-siRNA complexes was made in Optimem (Invitrogen) and mixed with 0.4μL Lipofectamine 2000 in 2.1μL Optimem for a total volume of 5μL per well, master mix is made based on how many wells needed. 1.25pmol transfection mix is made by diluting 2.5pmol transfection mix in equal value of Optimem, and 0.63pmol transfection mix is made by diluting 1.25pmol transfection mix in equal value of Optimem. Each transfection condition was prepared at least in triplicate. After a 24-h incubation, media in each well was removed, and luciferase activities were measured using Dual-Glo® Luciferase Assay System (Promega Corporation, Cat #E2920) based on manufacturer’s description.

**Fig. S1.**
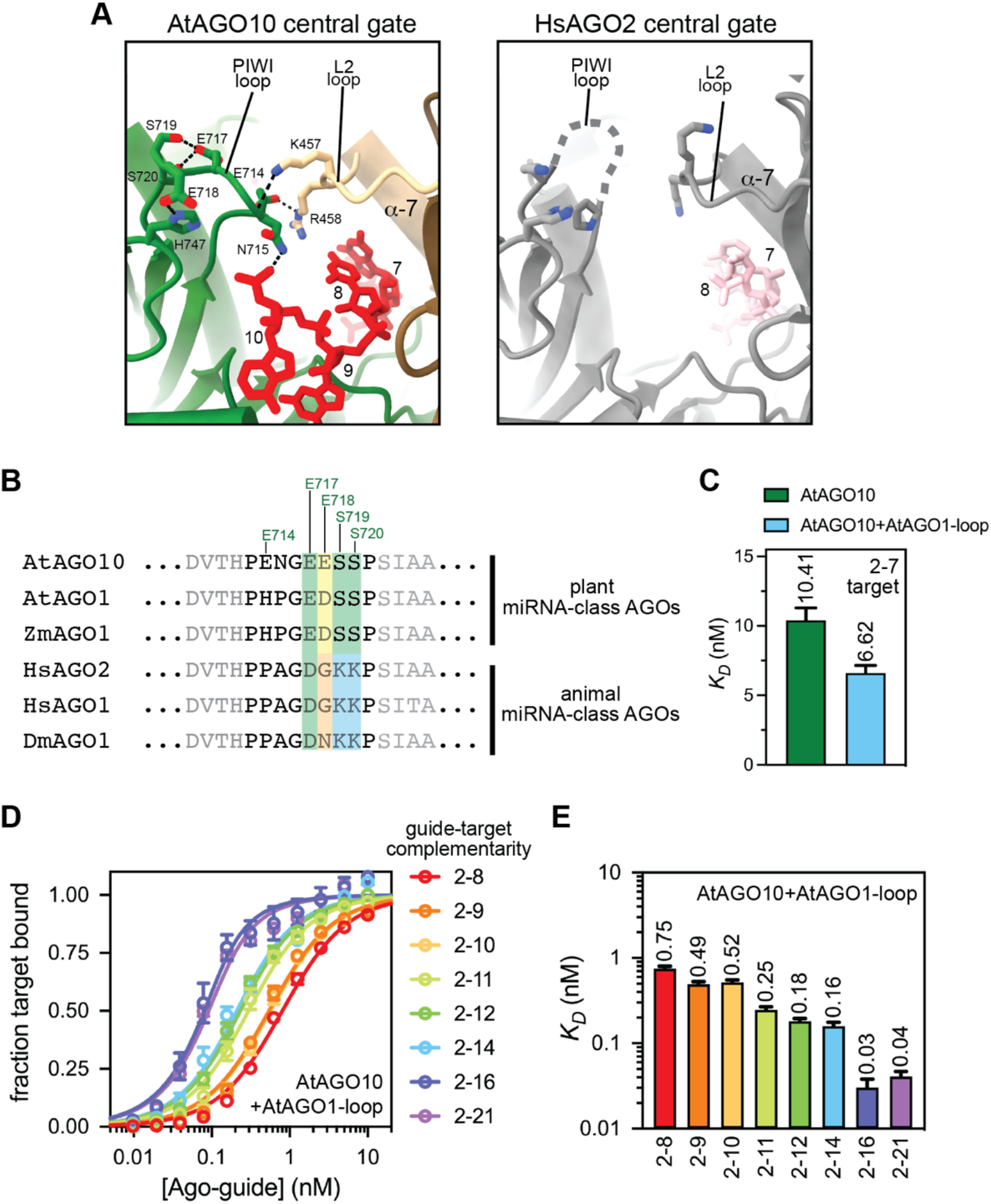
Supporting data for Fig. 1 and 2. **A**. Close-up views of PIWI-loop and L2-loop connections AtAGO10 and HsAGO2. **B**. Sequence alignment showing conservation of PIWI-loop sequence in plant clade I AGOs. **C**. Dissociation constants for wild-type AtAGO10 and a mutant AtAGO10 bearing the PIWI-loop sequence from AtAGO1 (AtAGO10+AtAGO10-loop) binding to a target RNA with complementarity to the canonical miRNA seed region (2–7). **D**. Fraction target RNA bound versus [AGO-guide] for the AtAGO1+AtAGO1-loop mutant. Guide and target RNAs are as shown in Fig. 2A. **E**. Dissociation constants derived from data in panel D.

**Fig. S2.**
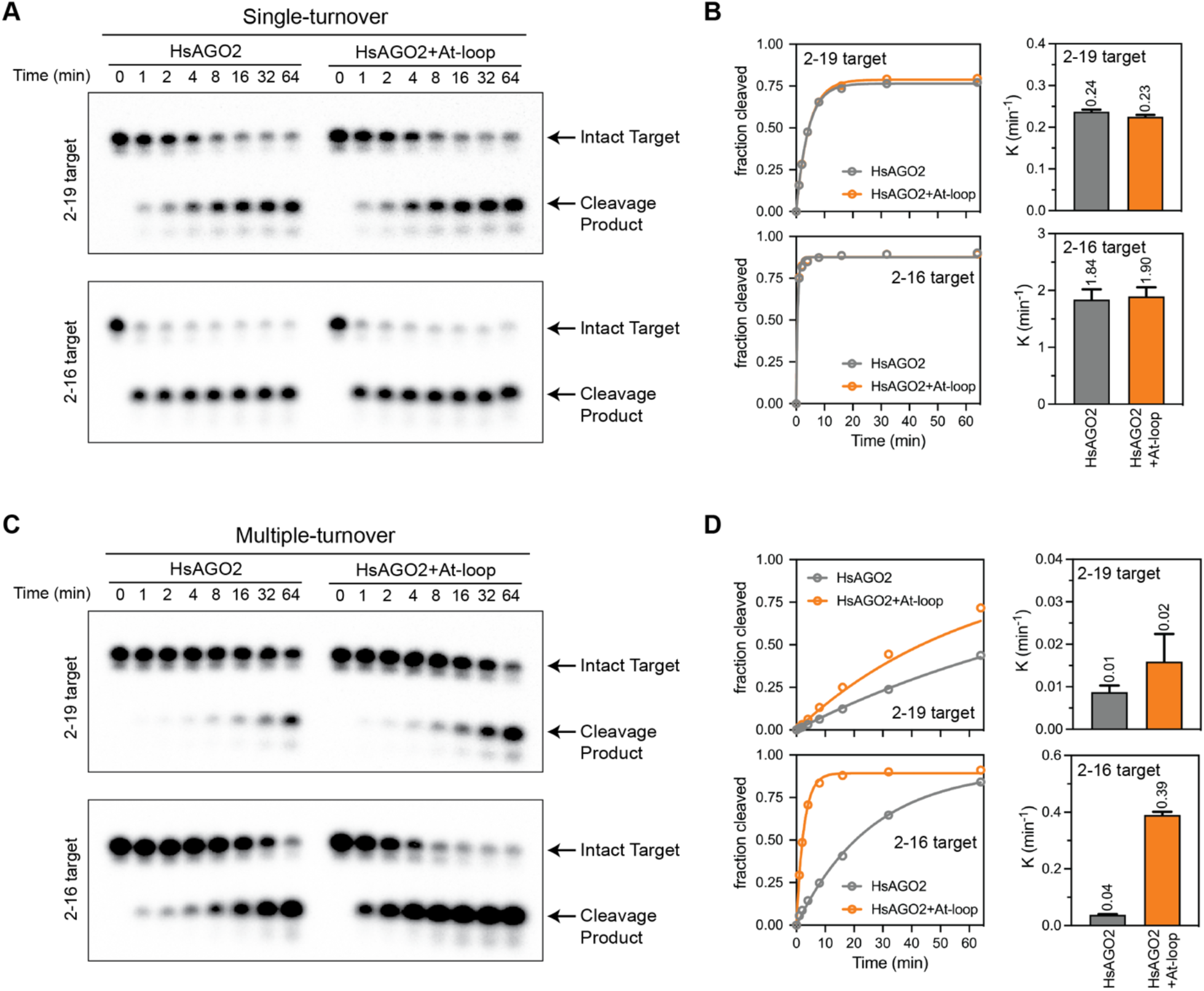
Additional target cleavage data supporting Fig. 4. **A**. Representative denaturing showing cleavage of 2–16 and 2–19 target RNAs by HsAGO2 and HsAGO2+At-loop under single turnover conditions after various amounts of time. **B**. Fraction of the target RNAs cleaved plotted versus time (left panel). The right panels show first-order rate constants for the single-turnover condition. **C**. Representative denaturing showing cleavage of 2–16 and 2–19 target RNAs by HsAGO2 and HsAGO2+At-loop under multiple turnover conditions after various amounts of time. **D**. Fraction of the target RNAs cleaved plotted versus time (left panel). The right panels show first-order rate constants for the multiple-turnover condition.

**Table S1.** Crystallographic and Refinement Statistics for human Ago2-miR-122 and human Ago2+At-loop-miR122 Complexes. Numbers in parentheses represent statistics in the highest resolution shell. R_merge_ = (Σ_h_Σ_i_|I_h_ - I_hi_|/Σ_h_Σ_i_I_h,i_) × 100, where I_h_ is the mean of I_h,i_ observations of reflection h. R-pim = Σ_h_ [1/(/n_h_ - 1)]^1/2^ Σ_i_|<I_h_> I_h,i_|/Σ_h_ Σ_i_ I_h,i_ R-work and R-free = ΣΠF_o_| - |F_c_Π/Σ|F_o_| × 100 for 95% of recorded data (R-work) or 5% of data (R-free). RMSD, root-mean-square deviation. CC, correlation coefficient.

| Structure | HsAGO2-miR122 | HsAGO2+At-loop-miR122 |
| --- | --- | --- |
| PDB Code | 8D71 | 8D6J |
| Wavelength (Å) | 0.97946 | 0.97946 |
| Resolution range (Å) | 37.71 - 2.5 (2.589 - 2.5) | 37.69 - 2.5 (2.589 - 2.5) |
| Space group | P 1 21 1 | P 1 21 1 |
| Unit cell parameters | 63.13 Å, 107.74 Å, 68.82 Å,<br>90°, 106.89°, 90° | 63.30 Å, 107.51 Å, 68.71 Å,<br>90°, 106.90°, 90° |
| Total reflections | 60147 (5968) | 60166 (5980) |
| Unique reflections | 30149 (2990) | 30141 (2992) |
| Multiplicity | 2.0 (2.0) | 2.0 (2.0) |
| Completeness (%) | 98.72 (99.27) | 98.75 (99.17) |
| Mean I/sigma(I) | 23.81 (4.81) | 12.90 (2.58) |
| Wilson B-factor | 47.93 | 49.32 |
| R-merge | 0.01911 (0.1429) | 0.03704 (0.2961) |
| R-meas | 0.02702 (0.2021) | 0.05239 (0.4187) |
| R-pim | 0.01911 (0.1429) | 0.03704 (0.2961) |
| CC1/2 | 0.999 (0.956) | 0.998 (0.832) |
| CC* | 1 (0.989) | 0.999 (0.953) |
| Reflections used in refinement | 30144 (2990) | 30135 (2992) |
| Reflections used for R-free | 1224 (122) | 1313 (134) |
| R-work | 0.2242 (0.2756) | 0.2326 (0.2996) |
| R-free | 0.2800 (0.3314) | 0.2854 (0.3359) |
| CC (work) | 0.942 (0.856) | 0.937 (0.802) |
| CC (free) | 0.897 (0.796) | 0.871 (0.691) |
| Number of non-hydrogen atoms | 6780 | 6774 |
| macromolecules | 6716 | 6723 |
| ligands | 1 | 1 |
| Protein residues | 810 | 810 |
| RMSD (bonds) | 0.006 | 0.004 |
| RMSD (angles) | 0.61 | 0.58 |
| Ramachandran favored (%) | 94 | 93.86 |
| Ramachandran allowed (%) | 5.38 | 6.02 |
| Ramachandran outliers (%) | 0.62 | 0.13 |
| Rotamer outliers (%) | 0.97 | 1.11 |
| Clashscore | 4.86 | 7.1 |
| Average B-factor | 60.7 | 59.26 |
| macromolecules | 61.13 | 59.59 |
| ligands | 49.71 | 50.11 |

